# Investigating the impact of database choice on the accuracy of metagenomic read classification for the rumen microbiome

**DOI:** 10.1101/2022.04.26.489553

**Authors:** Rebecca H. Smith, Laura Glendinning, Alan W. Walker, Mick Watson

**Author notes:** Corresponding author: Rebecca H. Smith.

## Abstract

Microbiome analysis is quickly moving towards high-throughput methods such as metagenomic sequencing. Accurate taxonomic classification of metagenomic data relies on reference sequence databases, and their associated taxonomy. However, for understudied environments such as the rumen microbiome many sequences will be derived from novel or uncultured microbes that are not present in reference databases. As a result, taxonomic classification of metagenomic data from understudied environments may be inaccurate. To assess the accuracy of taxonomic read classification, this study classified metagenomic data that had been simulated from cultured rumen microbial genomes from the Hungate collection. To assess the impact of reference databases on the accuracy of taxonomic classification, the data was classified with Kraken 2 using several reference databases. We found that the choice and composition of reference database significantly impacted on taxonomic classification results, and accuracy. In particular, NCBI RefSeq proved to be a poor choice of database. Our results indicate that inaccurate read classification is likely to be a significant problem, affecting all studies that use insufficient reference databases. We observe that adding cultured reference genomes from the rumen to the reference database greatly improves classification rate and accuracy. We also demonstrate that metagenome-assembled genomes (MAGs) have the potential to further enhance classification accuracy by representing uncultivated microbes, sequences of which would otherwise be unclassified or incorrectly classified. However, classification accuracy was strongly dependent on the taxonomic labels assigned to these MAGs. We therefore highlight the importance of accurate reference taxonomic information and suggest that, with formal taxonomic lineages, MAGs have the potential to improve classification rate and accuracy, particularly in environments such as the rumen that are understudied or contain many novel genomes.

## Background

Ruminants are vital for global food security, providing high-quality protein to the increasing food demands of an expanding human population. The rumen is home to a complex microbial ecosystem containing bacteria, archaea, fungi, protozoa and viruses. The relationship between the host and these microbes is symbiotic, as they ferment lignocellulosic feed into volatile fatty acids, which are a key energy source for the host animal [1]. Subsequently the rumen microbiome significantly contributes to global food security and world trade. Cows alone contribute substantially to the economy; in 2018 the global production value of beef exceeded $220 trillion USD, and cow’s milk exceeded $288 trillion USD (FAOSTAT). Understanding the rumen is paramount to the success of many avenues of agricultural research, including feed-conversion efficiency [2], [3], methane emissions [4–7] and investigating the impact of diet on antibiotic resistance [8].

In spite of the importance of ruminants, the rumen continues to be an under-characterised environment [9] with many ruminant-dwelling microbes remaining uncultured, and as such absent from public reference databases. To mitigate this issue, efforts have been made to culture ruminant-dwelling microbes, such as the Hungate 1000 project. This significantly improved knowledge surrounding ruminant microbiome community structure as these cultured microbes are estimated to represent approximately 75% of ruminant genera [10]. However, while culturing efforts have undoubtedly improved the availability of rumen isolated genomes, culturing is laborious, and some species may prove difficult to isolate in the laboratory. As a result, it is known that many ruminant genera remain to be cultured, and are therefore without sequence information [11], meaning reference databases still have important limitations.

Metagenomics is the simultaneous study of DNA extracted from organisms within an environment or microbiome (reviewed in [12]). Metagenome-assembled genomes (MAGs) are draft genomes that have been assembled ‘*de novo’*, without a reference genome, from binning metagenomic sequencing data [13]. As this process does not require culturing, MAGs can considerably expand on the number of reference genomes derived from culture collections. Additionally, MAG assembly is high throughput, hundreds or thousands of MAGs can be assembled during a single analysis. MAGs therefore have the potential to transform microbiome analysis by shedding light on the previously described “uncultured majority” ([14], [15]) and a recent cross-study examination of over 33,000 rumen MAGs concludes that there are still more ruminant species to discover [16]. As the rumen microbiome still remains predominantly uncultivated, the use of culture-independent techniques such as MAG assembly are therefore becoming increasingly valuable. Many novel MAGs have been recently published from ruminants [13, 17–25], and these allow the discovery of novel putative genes and functionality [26–28].

Studying the microbial composition of an environment using metagenomic data, necessitates the assignment of taxonomic labels to sequence reads, referred to as taxonomic read classification. Classification can be to varying taxonomic levels or ranks. Two of the most commonly used bioinformatics tools available for metagenomic read classification are Kraken [29], and its successor, Kraken 2 [30]. Regardless of classification tool used, reference database quality and comprehensiveness fundamentally underpin the accuracy of results, and classification results can vary dramatically depending on which reference database is used. However, reference databases are known to be highly skewed towards certain well studied species. Blackwell *et al.* showed that 90% of microbial genomes in the European Nucleotide Archive (ENA), a large publicly available microbial sequence archive, originate from just 20 species [31]. This is important because Meric *et al.* demonstrated that the number of genomes used to build the index, and the taxonomic system used to classify genomes, can significantly impact classification rates [32]. Similarly, Nasko *et al.* demonstrated that classification accuracy is impacted by the version of the popular publicly available sequence database [33] RefSeq that is used [34], and Marcelino *et al.* showed that the reference database needs to represent all domains of life within the microbiome to minimise false positives [35]. Of note, some rumen metagenomics studies report very poor read classification rates when using RefSeq alone [13], [17]. The Hungate 1000 project provides excellent additional reference genomes for taxonomic classification [10] but, given that there are hundreds of currently uncultured and uncharacterised genera in the rumen, the Hungate collection alone may not be fully representative. Subsequently, although the Hungate genomes may improve the classification rate of metagenomic data [13], these may not be true hits, and therefore may not always improve the accuracy of classification. Stewart *et al.* have twice demonstrated that the addition of MAGs to reference databases improves metagenomic read classification rate by 50-70%, but the addition of Hungate collection genomes showed little improvement (10%) [13], [17]. However, the impact of the addition of MAGs and Hungate collection genomes to reference databases on classification accuracy, not just classification rate, is not yet known.

In this study, simulated data generated from known rumen microbe genomes, was used to test the accuracy of metagenomic read classification using a range of reference databases. This work focused on the read classification tool, Kraken2, which has been shown to be highly accurate and fast [36] and allows for the easy construction of custom reference databases. We found that classification accuracy varies significantly between reference databases, and taxonomic levels. This work emphasises the importance of reference database choice, as well as highlighting the potential low accuracy of taxonomic classification using commonly-applied present approaches. Furthermore, this study demonstrates that the addition of MAGs to reference databases substantially improves read classification accuracy at some taxonomic levels. This work proposes that this improvement has the most potential when using MAGs assembled from the same environment as the classification data, and when using reference MAGs that have a full taxonomic lineage assigned to them.

## Results

### Classification rate is heavily impacted by reference database

In order to assess the impact of reference database choice on the classification of metagenomic data, a simulated metagenomic dataset was created from rumen microbial genomes. The taxonomy of the simulated metagenomic dataset was classified using Kraken2 and a variety of reference databases. Briefly, the ‘Hungate’ database contains rumen microbial genomes. The ‘RefSeq’ and ‘Mini’ databases contain the complete bacterial, archaeal and viral genomes in RefSeq, the human genome, as well as a collection of known vectors (UniVec_Core), with the ‘Mini’ database built to just 8 GB in size. The ‘RUG’ database contains rumen uncultured genomes (RUGs), which are MAGs that have been assembled from rumen metagenomic data. The ‘RefHun’ database contained the same sequences as the ‘RefSeq’ database, with the addition of the cultured isolate genome sequences in the ‘Hungate’ database. Similarly, the ‘RefRUG’ database contains the same sequences as the ‘RefSeq’ database, with the addition of the MAG sequences in the ‘RUG’ database. Further information on the contents of each database and how they were made can be found in the Methods section, and in Table 1.

**Table 1.** The contents of each reference database and instructions on how they were built. The six reference databases each contain different reference sequences, as described in the Table. Also shown are the commands used to download and/or add to the library for each database, and build each database using Kraken 2.

| Database | Contents | Construction |
| --- | --- | --- |
| Hungate | Custom database containing 460 rumen microbial reference genomes from the Hungate collection (see Supplementary Table S3) | `for file in /hungate_genomes/*.fasta do kraken2-build --add-to-library $file --db hungate_only_db_k2 done kraken2-build --build --threads 16 --db hungate_only_db_k2` |
| Mini | The complete collection of genomes in RefSeq for bacterial, viral and archaeal domains, the human genome and UniVec_Core vectors. The database was built to 8 GB in size to replicate the “MiniKraken” functionality of Kraken1 | `kraken2-build --download-library bacteria --db mini_standard_db_k2 --use-ftp kraken2-build --download-library archaea --db mini_standard_db_k2 --use-ftp kraken2-build --download-library viral --db mini_standard_db_k2 --use-ftp kraken2-build --download-library human --db mini_standard_db_k2 --use-ftp kraken2-build --download-library UniVec_Core - -db mini_standard_db_k2 --use-ftp kraken2-build --db mini_standard_db_k2 --build --max-db-size 8000000000 --threads 4` |
| RefSeq | The complete collection of genomes in RefSeq for bacterial, viral and archaeal domains, the human genome and UniVec_Core vectors | `kraken2-build --download-library bacteria --db standard_db_k2 --use-ftp kraken2-build --download-library archaea --db standard_db_k2 --use-ftp kraken2-build --download-library viral --db standard_db_k2 --use-ftp kraken2-build --download-library human --db standard_db_k2 --use-ftp kraken2-build --download-library UniVec_Core - -db standard_db_k2 --use-ftp kraken2-build --build --threads 16 --db standard_db_k2` |
| RUG | Custom database containing 4,941 rumen metagenome-assembled genomes (named “RUGs” - see Stewart *et al.* [17]) | `for file in /rug_drafts/*.fna do kraken2-build --add-to-library $file --db rug2_only_db_k2 done` |
|  |  | kraken2-build --build --threads 8 --db<br>rug2_only_db_k2 |
| RefRUG | The complete collection of genomes in RefSeq for bacterial, viral and archaeal domains, the human genome and UniVec_Core vectors with the addition of 4,941 rumen metagenome-assembled genomes (see RUG database) | kraken2-build --download-library bacteria --db<br>standard_rug2_db_k2 --use-ftp<br>kraken2-build --download-library archaea --db<br>standard_rug2_db_k2 --use-ftp<br>kraken2-build --download-library viral --db<br>standard_rug2_db_k2 --use-ftp<br>kraken2-build --download-library human --db<br>standard_rug2_db_k2 --use-ftp<br>kraken2-build --download-library UniVec_Core -<br>-db standard_rug2_db_k2 --use-ftp<br>for file in /rug_drafts/*.fna<br>do<br>kraken2-build --add-to-library \$file --db<br>standard_rug2_db_k2<br>done<br>kraken2-build --build --threads 16 --db<br>standard_rug2_db_k2 |
| RefHun | The complete collection of genomes in RefSeq for bacterial, viral and archaeal domains, the human genome and UniVec_Core vectors with the addition of 460 reference genomes from the Hungate collection (see Hungate database section of this table and Supplementary Table S3) | kraken2-build --download-library bacteria --db<br>standard_hungate_db_k2 --use-ftp<br>kraken2-build --download-library archaea --db<br>standard_hungate_db_k2 --use-ftp<br>kraken2-build --download-library viral --db<br>standard_hungate_db_k2 --use-ftp<br>kraken2-build --download-library human --db<br>standard_hungate_db_k2 --use-ftp<br>kraken2-build --download-library UniVec_Core -<br>-db standard_hungate_db_k2 --use-ftp<br>for file in /hungate_genomes/*.fasta<br>do<br>kraken2-build --add-to-library \$file --db<br>standard_hungate_db_k2<br>done<br>kraken2-build --build --threads 16 --db<br>standard_hungate_db_k2 |

As a first test, we looked simply at how much of the simulated metagenomic data was classified (classification rate), regardless of whether or not the classification was accurate. The overall classification rate, meaning the percentage of reads classified by Kraken2 to any taxonomic level when using that particular database, is shown in Figure 1. Also shown in Figure 1 is the percentage of reads that were unclassified by Kraken2, meaning they were not classified to any taxonomic level when using that particular database. As expected, since the simulated dataset was derived from the Hungate collection genomes, when the Hungate reference database was used Kraken2 classified almost all reads, with a classification rate of 99.95 %. The Kraken2 Mini and RefSeq reference databases resulted in the classification of 39.85 % and 50.28 % of the reads respectively. Interestingly, of the 460 Hungate genomes used to create the simulated data, 119 were present in RefSeq at the time of analysis. However, as Kraken 2 chooses which genomes to include in each Standard database, not all 119 Hungate genomes in RefSeq were necessarily included in the RefSeq or Mini databases. This indicates that the RefSeq database is not fully representative of the data, which will have impacted on the classification results. The RUG reference database alone had a classification rate of 45.66 %, which is a higher rate than the Mini Kraken 2 database but lower than the RefSeq database. Adding the RUG data to the RefSeq database (RefRUG) resulted in 70.09 % of reads being classified, which is approximately 1.4x as many reads than were classified with the RefSeq database alone. Finally, as expected, adding the Hungate database to the RefSeq database (RefHun) resulted in near complete classification of the reads. However, there was no apparent benefit to classification rate with the addition of RefSeq (RefHun), when compared to the Hungate database alone (Figure 1).

**Figure 1.**
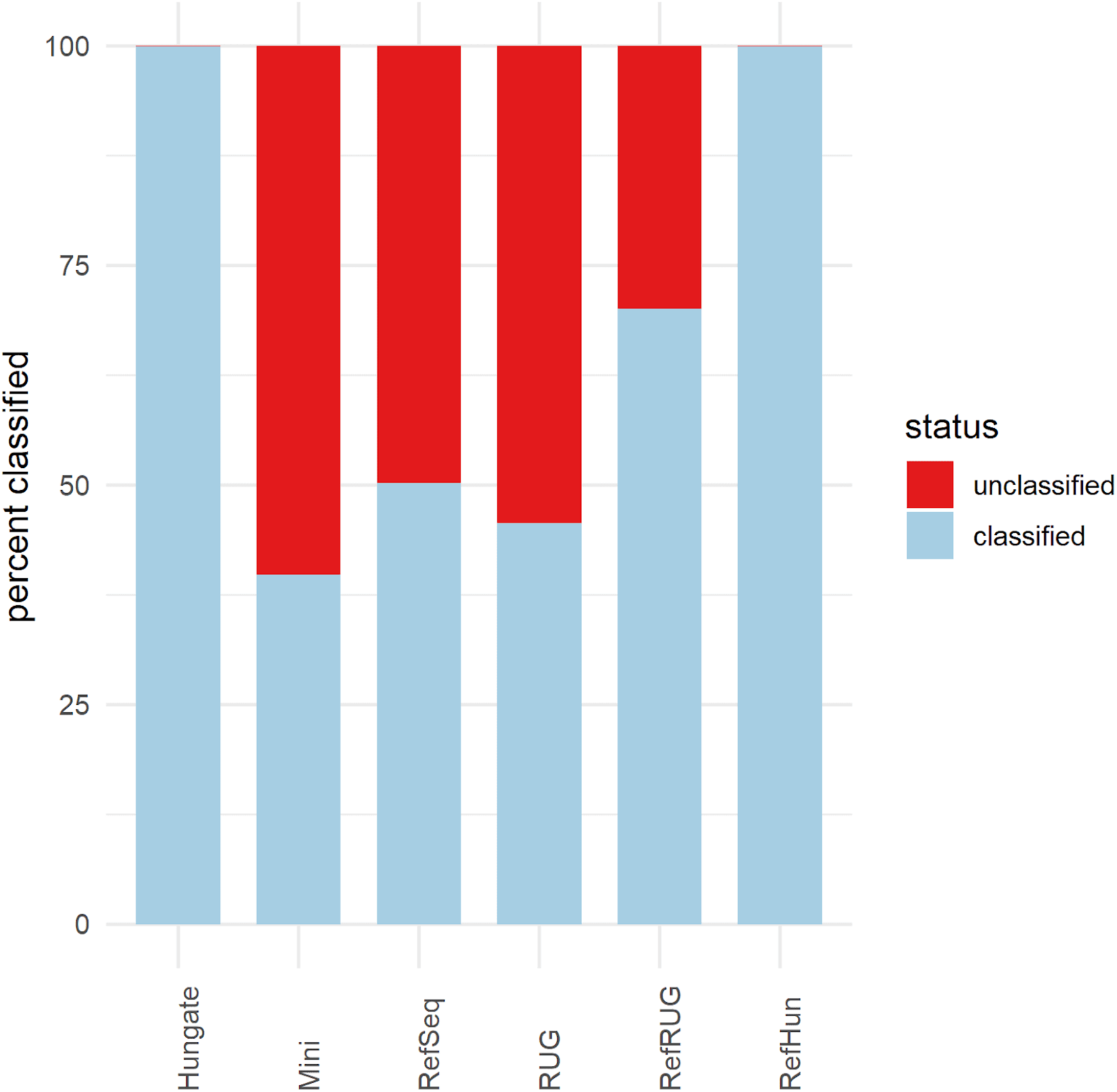
Overall classification rate of reads for the six reference databases. The classification rate of the data for each database are shown in the bars along the x-axis. Details about the databases can be found in Table 1. The y-axis denotes the percentage of reads from the simulated metagenomic dataset which were classified or unclassified by Kraken2 to any taxonomy level using each reference database.

After observing the overall classification rates for each reference database, the next step was to examine the classification rates at various taxonomic levels for each reference database. Figure 2 separates the overall classification rate for each reference database into the classification rate at various taxonomic levels. Overall classification rates, regardless of accuracy, are also shown in Supplementary Table S1. In general, there was a decline in the classification rate for each database moving down the taxonomic levels from phylum, to family, to genus and finally species.

**Figure 2.**
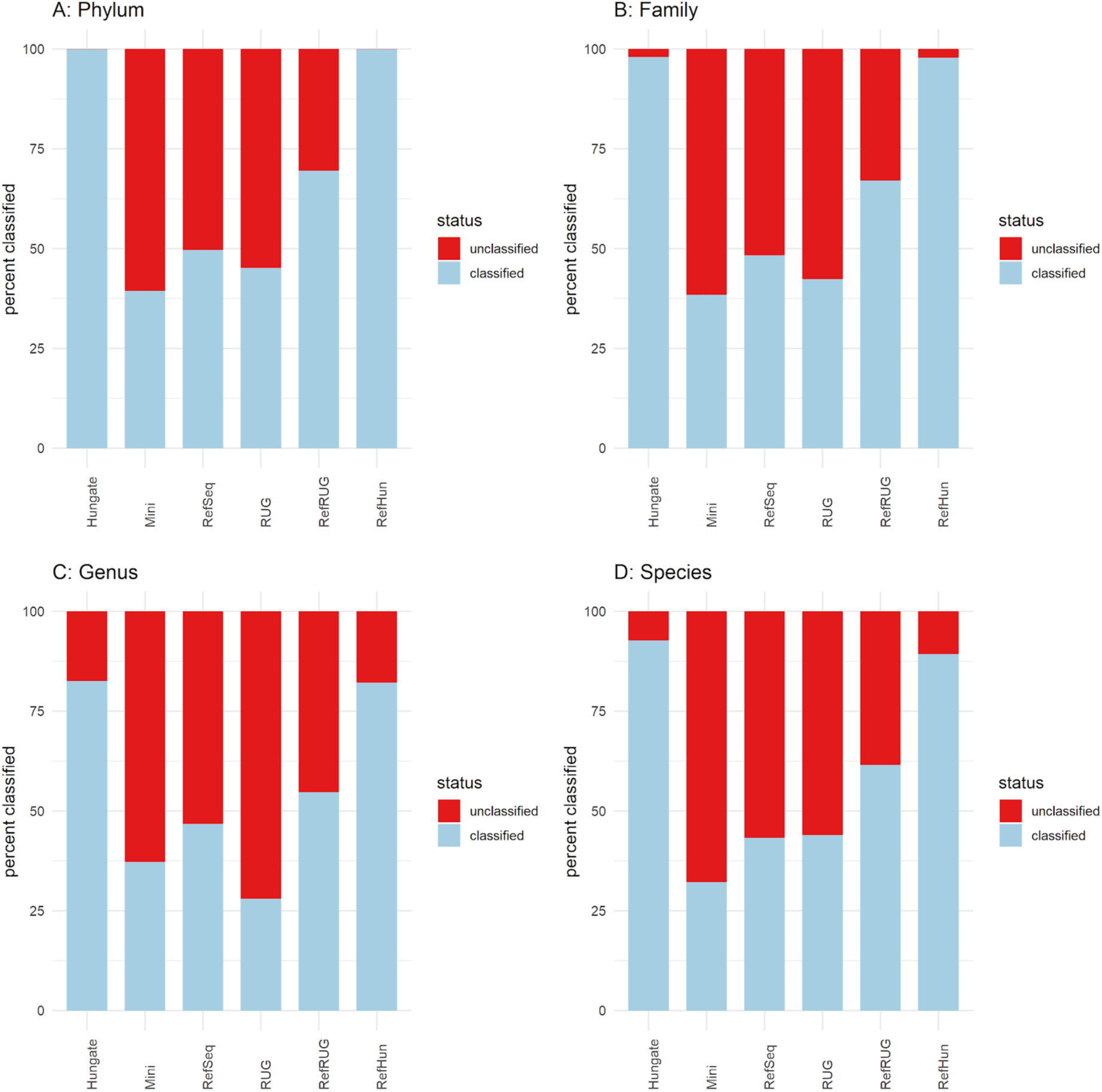
Classification rate of reads, shown at various taxonomic levels for the six reference databases. Classification rate refers to whether the reads were classified or unclassified, and are shown as a percentage at the (A) Phylum, (B) Family, (C) Genus and (D) Species levels. The y-axis shows the percentage of reads from the simulated dataset which were classified or unclassified when classified using Kraken2. The six reference databases used during classification are shown as bars plotted along the x-axis.

Anomalously, with some reference databases, classification rate at the genus level was lower than at the species level. This was also observed to a lesser extent in the classification rates at the family level. For example, the RUG database had a classification rate of 45.16% at phylum level, 42.36% at family level, 27.99% at genus level and 43.93% at species level. This is due to a feature of the data itself, as some of the Hungate and RUG genomes used to build the reference databases do not have complete taxonomic lineages. For example, the Hungate genome “*Bacteroidales* bacterium KHT7” (taxonomy ID: 1855373) has labels at the kingdom, phylum, class, order and species levels, but no labels at the family and genus levels. Of the 460 Hungate genomes, 8 do not have a label at the family level, and 73 do not have a label at the genus level. Another example is the RUG “*Ruminococcaceae* bacterium RUG10048” (taxonomy ID: 1898205), which has the label *Ruminococcaceae* at the family level, and the label “*Ruminococcaceae* bacterium” at the species level, but has no label at the genus level. Of the 4941 RUGs, 3849 have no labels at the genus level, and 1753 have no labels at the family level. 4293 of the RUGs had a non-specific species label, for example “uncultured *Bifidobacterium* sp.”. Therefore, as these genomes do not have a taxonomic label at these levels, reads from these genomes appear as unclassified.

The addition of RefSeq to the Hungate reference database (RefHun database) did not significantly impact the classification rate at the higher taxonomic levels compared to the Hungate reference alone (Figure 2). However, at the lower taxonomic levels, the RefHun database appeared to slightly reduce the classification rate when compared to the Hungate database alone. For example, at the species level with the Hungate database 92.69% of reads were classified, whereas with the RefHun database 89.27% of reads were classified.

### Classification accuracy is strongly impacted by reference database

Although classification rate is an important feature, it is clearly more important that data that is classified is done so accurately. The next logical step was therefore to use ground truth data to investigate the read classification accuracy of each reference database on the simulated metagenomic data. Figure 3 shows the classification accuracy of reads when classified using each reference database, at various taxonomic levels. The same data in tabular form is shown in Supplementary Table S2. The percentage of correctly classified reads reduced when moving down the taxonomic levels from phylum to species, for all databases. At the phylum level, the majority of taxonomic labels assigned to classified reads were correct when using all reference databases, or were otherwise unclassified. Indeed, fewer than 4% of classified reads were classified incorrectly for any of the databases at the phylum level.

**Figure 3.**
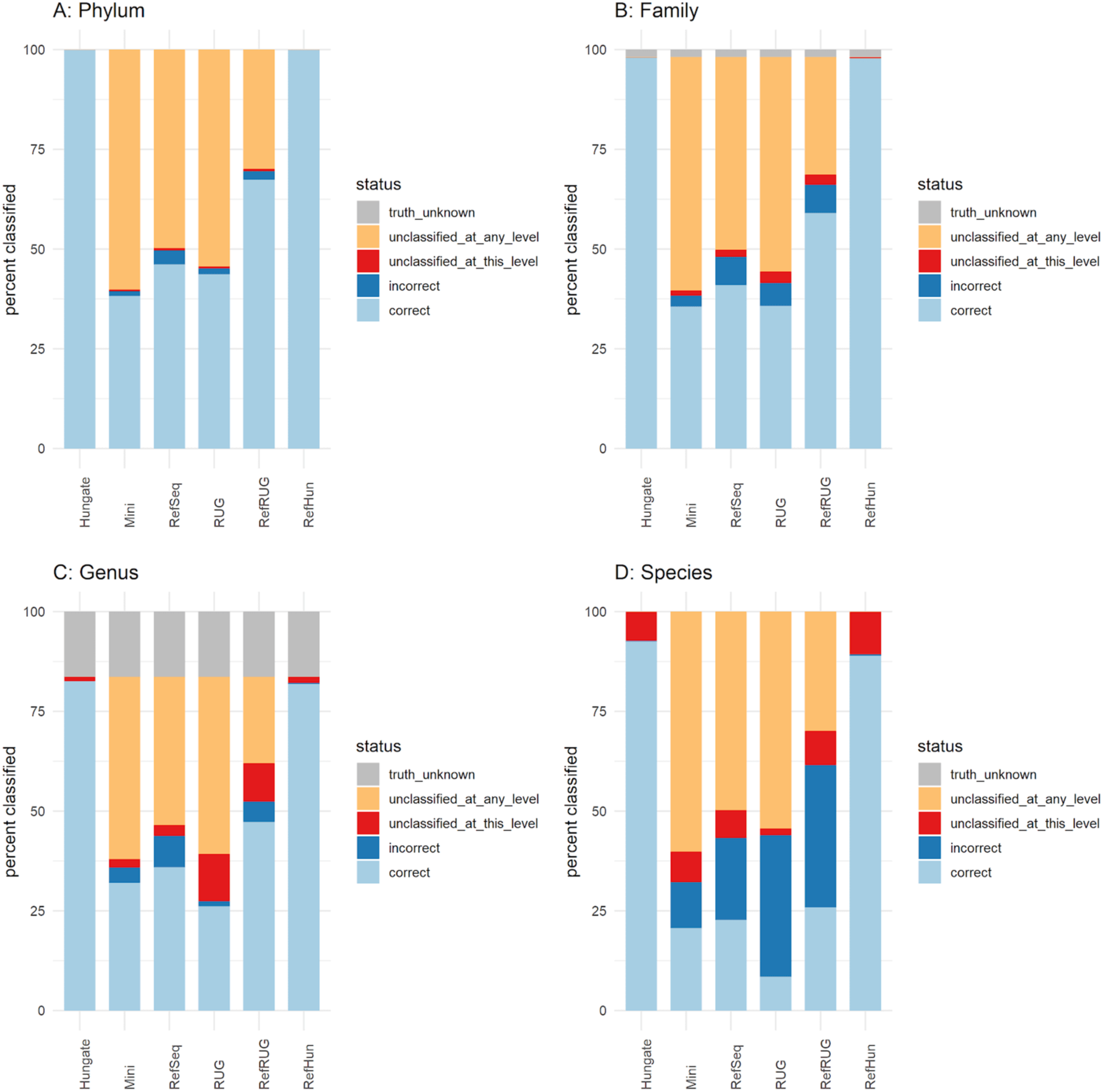
The accuracy of taxonomic classification using each reference database and across the various taxonomic levels. Classification status of reads compared to the ground truth for the six reference databases at various taxonomic levels. The graphs refer to the percentage of reads, shown along the y-axis, at the (A) Phylum, (B) Family, (C) Genus and (D) Species levels. Each bar represents reads classified by Kraken2, using each reference database as shown along the x-axis. The bars represent the percentage of classified reads at various classification status, as shown in the key. “Truth unknown” refers to the reads that originate from genomes that do not have an assigned family or genus. “Unclassified at any level” refers to reads that were not classified to any taxonomic level. “Unclassified at this level” refers to reads that were classified at other taxonomic levels, but not the level being examined in each graph. “Correct” and “incorrect” refer to reads that were classified correctly or incorrectly by Kraken2 using the respective database.

At the family level and above, no reads were classified incorrectly by Kraken2 with the Hungate database. The addition of Hungate genomes to the RefSeq database (RefHun) also increased the percentage of correctly classified reads substantially compared with using the RefSeq database alone, from 40.93% to 97.82%. Use of some of the reference databases resulted in reads being incorrectly classified at the family level. While classification using the RefSeq database correctly classified a higher percentage of reads than the Mini database (40.93% vs 35.62%), it also incorrectly classified a higher percentage (7.07% vs 2.74%), and the ratio of correct:incorrect was better when using the Mini database. Classification using the RUG database resulted in 35.76% of reads being classified correctly, which was less accurate than the RefSeq database but comparable to the Mini database. Additionally, use of the RUG database classified 5.71% of reads incorrectly, which was lower than the RefSeq database but higher than the Mini database. Adding the RUG genomes to the RefSeq database (RefRUG) improved almost all classification metrics when compared to using RefSeq alone. However, use of the RefRUG database resulted in a higher number of reads that were classified incorrectly (Figure 3). Use of the Hungate database correctly classified 97.99% of reads, and the remaining 2.01% were either unclassified or do not have a known truth due to missing taxonomic labels in the reference sequences. These reads are assigned the “truth_unknown” status.

At the genus level, although using the RefSeq reference database resulted in more reads being classified correctly than with the Mini database, using the RefSeq database also classified more reads incorrectly, with use of the Mini database again having a better ratio of correct:incorrect assignments. Using the RUG database resulted in fewer reads being classified correctly at the genus level, and resulted in a higher percentage of unclassified reads. However, use of the RUG database again resulted in fewer reads being incorrectly classified than with the RefSeq database. Similar to the family level results, adding the RUG data to RefSeq improved on most metrics when compared to using only the RefSeq database. Use of the Hungate database correctly classified 82.56% of reads, notably caused by reads categorised into the previously mentioned “truth_unknown” status, which accounted for 16.32% of the reads at genus level. Use of the Hungate database resulted in the incorrect classification of very few reads, which was echoed in the RefHun database. Compared to the RefSeq database, classification with the RefHun database classified more reads correctly (81.90% vs 35.97%), and classified fewer reads incorrectly (0.01% vs 7.85%).

At the species level, use of both of the RefSeq and the Mini databases classified a similar proportion of reads correctly (22.74% vs 20.65%). However, using the RefSeq database incorrectly classified almost the same proportion (20.53%), whereas using the Mini database incorrectly classified approximately half that amount (11.55%). As expected for a smaller database, classification with the Mini database had a higher proportion of reads that were unclassified at any level compared to RefSeq (60.15% vs 49.72%). A summary of the number of genera and species in the ground truth data, and the number that were classified using each of the reference databases, is shown in Supplementary Figure S1. Reference databases that include RefSeq (RefSeq, Mini, RefHun, RefRUG) classified thousands more false positives than databases that did not (Hungate, RUG). Including RUGs in the database (RUG) did not improve the situation, as it failed to classify many genera and species that were in the ground truth data. Additionally, classification of the data using the RUG database failed to classify any reads for certain abundant taxa.

After some investigation, it was discovered that there were marked differences in the annotated taxonomies present in the RUG and Hungate genomes, shown in Table 2. Several taxa were present in the Hungate data but were seemingly not present in the RUG data. As the Hungate collection contains highly abundant rumen microbial genomes, it is likely that these taxa are also present in the assembled RUG genomes, but that their taxonomy is not accurately annotated. Further investigation revealed that this was indeed a result of some RUGs not having an assigned taxonomy at the family and/or genus levels. Examples are the family *Bacteroidaceae* and genus *Bacteroides*, which are both present in the Hungate data but not annotated as such in the RUG data, explaining why no reads were classified for these taxa at those levels.

**Table 2.** The frequency of families and genera in the Hungate and RUG datasets, and overlap between the two datasets. Shown are the families and genera present in the Hungate and RUG datasets, including overlapping taxa. The Hungate data was used to generate the simulated data, and was included in the Hungate and RefHun reference databases. Similarly, the RUG data was included in the RefRUG and RUG reference databases.

| Status | Family | Genus |
| --- | --- | --- |
| Present in Hungate but not RUG | 25 | 48 |
| Present in RUG but not Hungate | 8 | 8 |
| Present in both RUG and Hungate | 23 | 33 |

The poor performance of RUGs at this level, as demonstrated in classification accuracy for the RUG database, also impacted the RefRUG database. Use of both reference databases including RUGs resulted in over 35% of reads being incorrectly classified. This can be explained by the use of generic species labels for the RUG dataset, which when compared to the formally named Hungate collection genomes in the ground truth were classified as incorrect. The addition of the RUG genomes to the RefSeq database (RefRUG) increased the percentage of correctly classified reads slightly, from 22.74% to 25.87%.

Once more, using the Hungate reference database resulted in the best performance, with the vast majority of reads classified correctly (92.56%), and only a small proportion of misclassifications (0.13%). There were, however, approximately 7% of reads that were not classified at the species level. The classification metrics when using the RefHun reference database were markedly closer to the results obtained when using the Hungate database than the RefSeq database. The addition of the Hungate genomes to the RefSeq database (RefHun) increased the percentage of correctly classified reads from 22.74% to 88.92%, and the decreased number of incorrectly classified reads from 20.53% to 0.35%, clearly demonstrating the huge gains in accuracy that can be obtained when closely matching sequences are present in reference databases.

### Composition of the reference database used impacts upon the accuracy of taxonomic read classification and taxonomic read abundance

Having demonstrated that the accuracy of taxonomic read classification changes considerably depending on the reference database used, this study next examined the impact of reference database choice on the taxonomic abundance of a microbial community. This was done using the same simulated data and reference databases as before, but by examining classification results in the form of taxonomic read abundance. Figure 4 shows a selection of scatterplots that compare the taxonomic abundance of the ground truth simulated metagenomic data with that of the classified data. The closeness-of-fit of the taxonomic read abundance (Figure 4) to the linear regression was measured using the R^2^ statistic, and is shown in Figure 5. The R^2^ statistic summarises how similar the classified taxonomic abundance was to the taxonomic abundance of the ground truth simulated data, and is therefore another indication of classification accuracy using each of the reference databases at various taxonomic levels.

**Figure 4.**
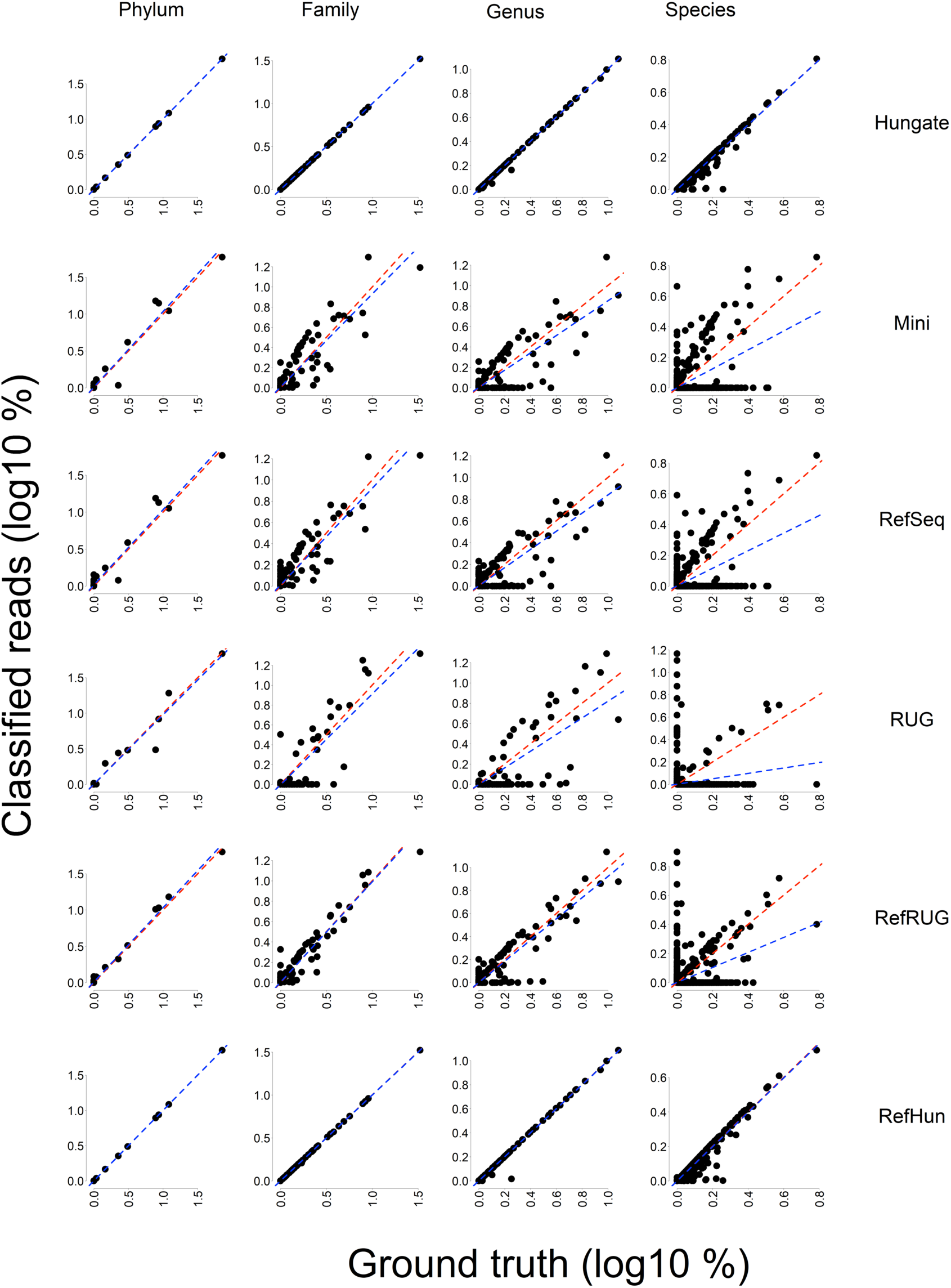
Comparing taxonomic abundance of the ground truth metagenomic data with that of the classified data. Scatterplots show the comparison between the simulated metagenomic data (ground truth, x-axis) and classified reads (y-axis). Data is plotted as a percentage of classified reads for the classified data, and a percentage of simulated reads for the ground-truth data. The data has been transformed by log10. A y=x line (shown in red) has been added to demonstrate how data points would appear on the graph if the number of ground-truth and classified reads were the same. A linear regression has been added (shown in blue) and used to calculate the R^2^ statistic, see Figure 6. Comparisons are shown at the Phylum, Family, Genus and Species levels, for the Hungate, Mini, RefSeq, RUG, RefRUG and RefHun reference databases.

**Figure 5.**
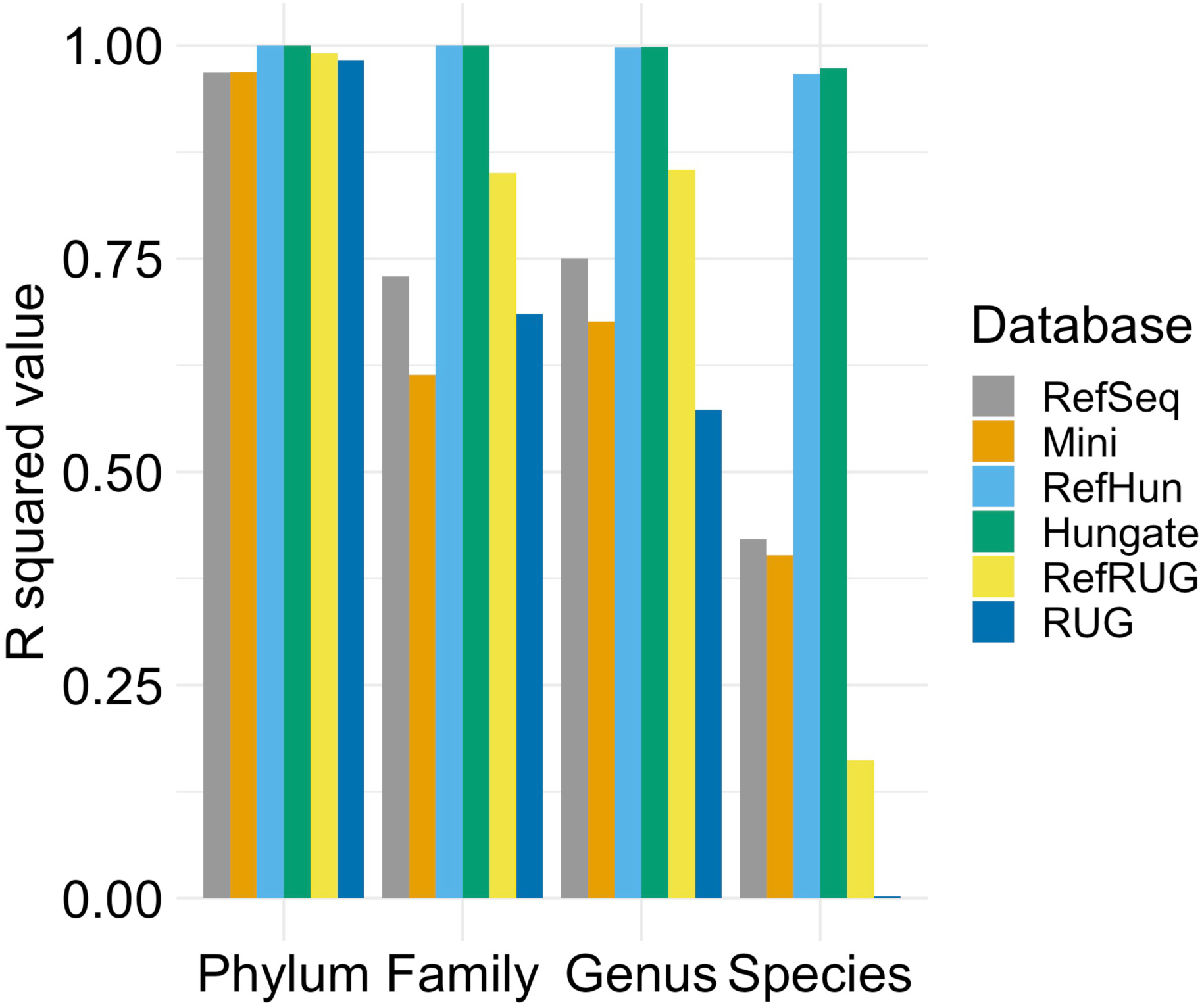
R^2^ values of the comparisons between taxonomy of the simulated metagenomic dataset and classified taxonomy at various taxonomic levels. The key denotes each reference database used to classify the data, and these are shown as individual bars at each taxonomic rank, displayed on the x-axis. The R^2^ value is the statistical measure of the correlation of data to the linear regression, measured using the scatterplots shown in Figure 4.

A cornerstone of microbiome research is community structure, which can be observed as a sample’s taxonomic abundance. To investigate this, the most abundant taxa in the ground truth data were observed in the classified data. Barplots displaying the taxonomic read abundance of the ground truth data, as well as the read abundance once the data was classified using each of the reference databases, are shown in Figure 6. Each plot shows the taxonomic distribution of the top 10 most abundant taxa for the ground truth data and the abundance of these taxa in the classified data, at that particular taxonomic level.

**Figure 6.**
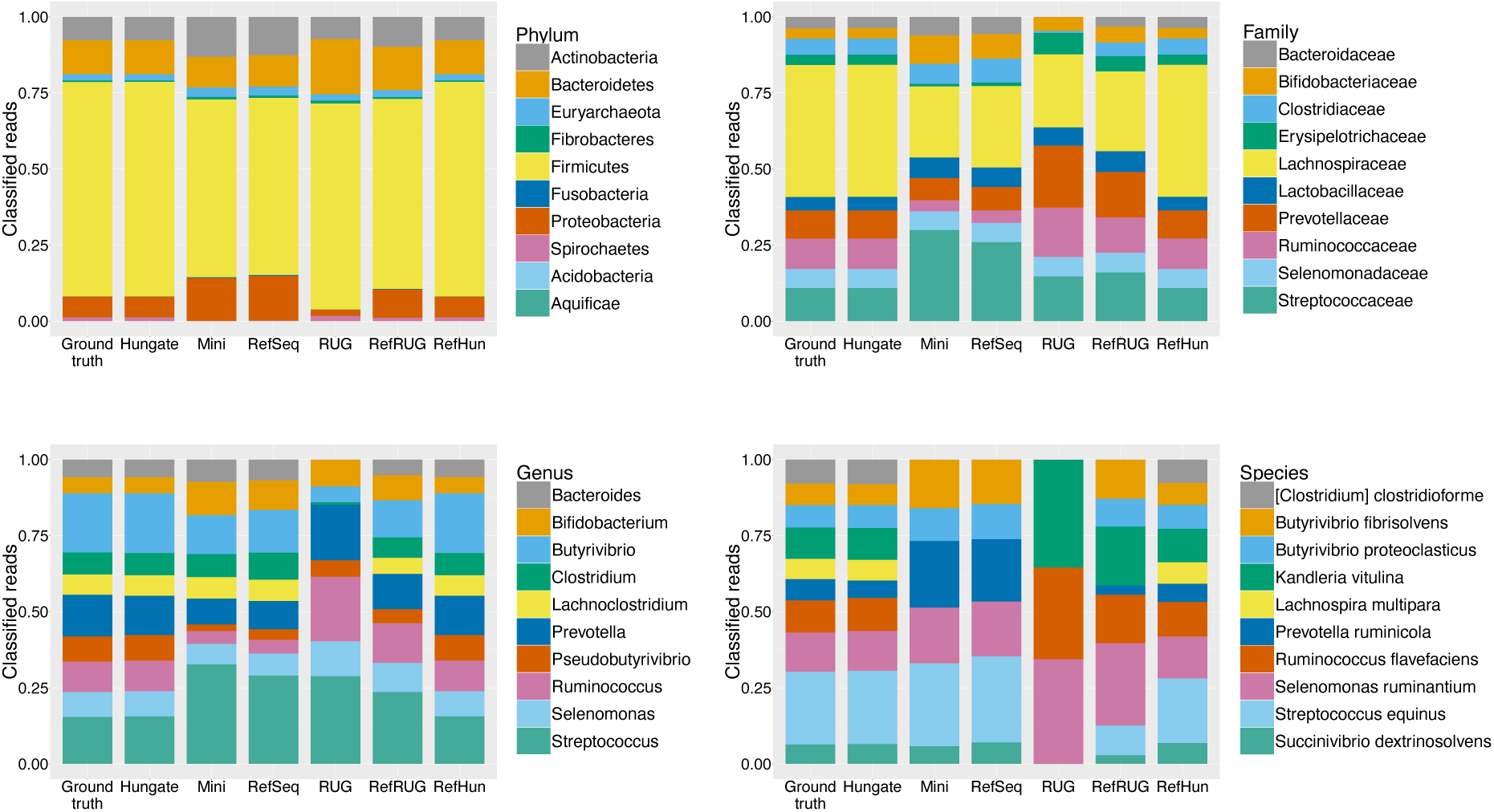
Comparing the classification of abundant taxa in the simulated metagenomic dataset for each reference database. Taxonomic distribution for the top ten most abundant taxa in the simulated metagenomic dataset, classified at the Phylum, Family, Genus and Species levels with Kraken2 using the six different reference databases. The y-axis denotes the percentage of reads classified at each level. The bars along the x-axis each represent the classification results for each database, split by taxonomy as shown in the keys for each level.

Overall, the Hungate and RefHun databases performed very well at classifying the data, as shown in Figures 4, 5 and 6. There was a slight reduction in accuracy at the species level, where the R^2^ value was 0.97, but this had little effect on the classification of abundant taxa (see Figure 6).

Using the RefSeq and Mini reference databases accurately classified the data at phylum level, but there was a distinct drop in accuracy at the class level, which continued further down the taxonomic levels. At the phylum level, the Mini and RefSeq databases over-estimated *Proteobacteria* and *Actinobacteria*, but under-estimated *Firmicutes*. At the family level, the Mini and RefSeq databases overestimated the *Streptococcaceae* and *Bifidobacteriaceae*, yet underestimated the *Lachnospiraceae* and *Erysipelotrichaceae*. At the genus level the Mini and RefSeq databases overestimated the *Streptococcus* and *Bifidobacterium*, and underestimated *Ruminococcus* and *Prevotella*. At the species level, the RefSeq and Mini databases did not classify any reads to four of the ten most abundant species: *Clostridium clostridioforme*, *Lachnospira multipara*, *Ruminococcus flavefaciens* or *Kandleria vitulina*.

The RUG and the RefRUG databases were similarly accurate at the phylum level, but began to diverge in classification accuracy at lower taxonomic levels. In general, the RefRUG database classified the data more accurately than the RUG database, and this was likely due to the issues surrounding taxonomic labelling of the RUGs, as described above. At the family level, the RUG database did not classify any reads as *Bacteroidaceae*, and at the genus level there were a lack of reads classified as *Bacteroides*. This was simply because these taxonomic labels do not appear in the RUG collection. At the species level, the RUG database classified just three of the top ten most abundant taxa in the simulated metagenome (Figure 6). This resulted in a poor correlation in Figure 4 and a very low R^2^ value of 0.002 (Figure 5). Interestingly, however, two out of the three species (*Ruminococcus flavefaciens* and *Kandleria vitulina*) were completely missed during classification by the RefSeq database, but were classified when the RUG data was added to the RefSeq database (RefRUG database). However, the species *Clostridium clostridioforme* and *Lachnospira multipara* were not classified when using the RefRUG reference database or indeed any databases other than Hungate or RefHun.

## Discussion

### Accuracy and rate of metagenomic data classification is heavily impacted by the choice of reference database

Research into microbiomes has increased substantially over the last two decades, driven by advances in DNA sequencing technologies. However, DNA-sequence based methods depend fundamentally on the quality of reference databases that are used to assign taxonomy or function to the sequence data. This study, which used a simulated metagenomic dataset, demonstrates the huge difference that choice of reference database can have on the accuracy of the results obtained.

RefSeq, the open-access database from NCBI, is a popular choice of reference database when classifying metagenomic data. However, using the RefSeq database we show that less than 40% of reads at genus level, and less than 25% of reads at species level, were accurately classified (Figure 3). Although this issue impacts all taxonomic levels, classification using these databases at the species level is particularly unreliable. When the data was classified using the RefSeq database, this study observes that nearly 50% of species taxonomy assignments were incorrect. This finding indicates that this frequency of inaccurate classification may be occurring in the many other studies that use the RefSeq database, compromising classification results. Use of the Mini database, which is optimised for use when there are limited computational resources available, also resulted in the classification of less than 40% of reads overall. This suggests that studies relying on the RefSeq or Mini database for classification will likely have a large proportion of inaccurate taxonomy assignments, which could impact strongly on subsequent interpretations and conclusions based on those results.

### Genomes from cultured isolates derived from the environment of study hugely increase classification rate and accuracy

Current reference databases are hugely biased towards microbes that have been isolated from well-studied environments, such as the 20 microbial species contributing to 90% of the reference genomes in the ENA [31]. The rumen is an under-studied environment, which has consequently impacted the number of ruminant microbial reference genomes present in public databases such as NCBI RefSeq. At the time of writing, of the 460 Hungate genomes used to create the simulated data, only 119 are present in NCBI RefSeq. The Kraken “Standard” database contains a subset of NCBI RefSeq, and so the RefSeq database may not contain all 119 of these Hungate genomes.

The Hungate reference database used here contained all of the Hungate genomes, and so is fully representative of the data that was classified. As expected, classification with the Hungate database resulted in classification of the majority of reads, and was the most accurate out of all the databases. However, at the species level, 7.31% of reads were not classified. Interestingly, these reads were unclassified rather than incorrectly classified. This reduction in classification at the species level was likely due to the phenomenon described by Nasko *et al.*: the so-called “minimiser collision”. This is where two distinct k-mers are minimised to identical minimisers (l-mers). In other words, if reads are highly similar, Kraken2 may be unable to distinguish between reference genomes at the species level, and so would assign taxonomy at the lowest common ancestor, therefore assigning taxonomy to a higher level [30].

In an attempt to understand the impact that including reference genomes from cultured representatives can have on classification accuracy of metagenomic data, we added the Hungate genomes to RefSeq, creating the RefHun reference database. Classification using the RefHun reference database showed significant improvements in classification rate and accuracy compared to the RefSeq database alone. This demonstrates that when classifying environmental data, classification accuracy can improve considerably by including more genomes derived from taxonomically well characterised cultured isolates in reference databases. Continued efforts to isolate, and formally taxonomically characterise, previously uncultured microbes from the rumen microbiome, and indeed any other understudied environment, is likely to have significant benefits for the accuracy of metagenomics-based studies.

### MAGs have the potential to improve metagenomic data classification even further, but are currently limited by their poorly defined taxonomy

While the addition of cultured isolate genomes clearly improves classification accuracy, it must be acknowledged that cultivation of microbes, and formally describing their taxonomy, are hugely time-consuming and labour-intensive activities [37]. Furthermore, many microbes may prove difficult to cultivate under laboratory conditions [38]. There are therefore significant bottlenecks that preclude the required widespread cultivation and characterisation of microbes. Therefore, the incorporation of MAGs, which can be generated without having to cultivate microbes in the laboratory, and can be done at far greater scale, in reference databases is an extremely promising additional or alternative avenue to improve classification of metagenomics datasets. In support of this, the addition of RUGs (MAGs) to the RefSeq database in this study (RefRUG) improved classification rate, which confirms the observations of other studies. Stewart *et al.* observed poor classification rates of rumen metagenomic data when using RefSeq, and reported the addition of Hungate collection genomes led to a classification rate increase of 2-fold, and the addition of RUGs led to an increase of 5-fold [13]. In a different study, Stewart *et al*. noted an increase of 10% in classification rate when adding Hungate collection genomes, and a 50-70% increase when adding RUGs to the reference database [17]. Xie *et al*. observed improvements in taxonomic classification rate with the addition of rumen MAGs to the reference database, compared with using Genbank and RMG entries alone [22].

Although addition of RUGs increased classification rate, using the RUG database resulted in the classification of reads with varying accuracy. In some respects, the effect was positive. For example, at the family and genus levels classification using the RUG database resulted in less reads being incorrectly classified than when using the RefSeq database. However, it is clear that there are likely to be significant issues with accuracy when using common current reference databases to classify metagenomic data. In this study, the ground truth information was available, which means we can say with certainty that some of the data was classified incorrectly. However, in real world scenarios, the correct taxonomy of the newly-sequenced data is of course unavailable, which means that the accuracy of classification results is difficult to quantify. We term such incorrectly classified reads as false positives, because in real world studies these incorrect classifications would be considered genuine. Marcelino *et al*. hypothesise that false positives occur as a result of conserved regions of reference genomes and sequence contamination in databases [35]. The use of each database classified some reads as false positives, although the highest number of false positives were classified by the reference databases containing RefSeq. In particular, classification using the RefSeq, Mini and RefRUG databases resulted in the apparent detection of thousands of species that were simply not there. The occurrence of false positives in this study indicates that false positives could be a common occurrence in metagenomic read classification.

More concerningly, addition of the RUG MAGs resulted in very poor overall classification accuracy, despite the addition of much more comprehensive reference material to the database. The likely explanation for this finding comes from the fact that, when the taxonomic labels in the Hungate and RUG data were compared at the family and genus levels, it was discovered that less than half of the total taxa were supposedly present in both datasets. As both data sets originate from the rumen, this is unlikely and is most probably a result of the incomplete and informal taxonomy labels used for the MAGs. This highlights the issue that reference sequences with incomplete or informal taxonomic labels may not be appropriate for classifying taxonomy. This issue can be resolved by ensuring all reference sequences, whether cultured isolate or MAG-derived, have complete, and accurate, labels across all taxonomic levels.

Taxonomy currently relies on consistent nomenclature to classify all organismal names across all living domains on Earth. NCBI taxonomy contained over 280,000 informal bacterial species (as of May 2017)[39], [40] and the NCBI databases contain 3760 genomes for unclassified or candidate bacteria at the time of writing. Issues arise when taxa are placed into a taxonomy database with informal names or incomplete lineages. For example, some of the Hungate collection genomes do not have an assigned rank at family or genus level. Additionally, assembled genomes (MAGs) often have an informal species name that does not follow traditional binomial nomenclature [41]. This issue was well demonstrated in this study, as classification using the RUG database failed to classify any reads from seven of the top 10 species in the ground truth data. This is surprising as these species are highly abundant in the rumen, and so you would expect to see them in the highly comprehensive RUG database. Of the 78 labels assigned at the species level by the RUG database, 56 had informal names, for example “uncultured *Lachnospiraceae* bacterium RUG10034”.

As MAGs are draft genomes, and can often be novel species or even novel clades, it can be difficult to correctly assign phylogeny and taxonomy. This is a significant problem, as metagenomics studies increasingly demonstrate that the rumen contains many genomes that cannot be easily placed into the current NCBI taxonomy. For example, Stewart *et al*. [17] found that of 4941 MAGs, 4303 could not be assigned a species, 3849 could not be assigned a genus, 1753 could not be assigned a family and 140 could not be assigned a phylum. However, this issue of uncertain phylogeny placement is not unique to MAGs, an example being the genus *Clostridium*, which has been demonstrated to actually consist of multiple genera [42]. Regardless of whether genomes are derived from cultured isolates or MAGs, mistakes or gaps in taxonomic descriptors will impact the accuracy of taxonomic classification.

It has been suggested that a change in microbial taxonomy towards a genome-based approach would improve upon the current taxonomy [43], [44]. The Genome Taxonomy Database (GTDB) uses a genome-based taxonomy, assigning the taxonomy of genomes based on their phylogeny [45]. Glendinning *et al*. observed many discrepancies between the phylogeny of MAGs and NCBI taxonomy, which was not found when using GTDB [24].

## Conclusions

In this study, we compare taxonomic classification results with ground truth simulated metagenomic data. Our results show that classification rate, classification accuracy and taxonomic read classification are heavily impacted by the choice of reference database used. In particular, RefSeq alone is a poor choice for classifying ruminant metagenomic data. Notably, our results indicate the extent to which ruminant metagenomic data could be inaccurately classified, an issue that has the potential to affect all studies that use insufficient reference databases. We demonstrate that custom reference databases substantially improve classification accuracy, and that genomes derived from cultured representatives and MAGs improve classification rate in all cases, but only improve classification accuracy for levels in which they have assigned taxonomy. This highlights the opportunity of using MAGs to improve taxonomic classification results in under-characterised environments, but also emphasises the importance of complete taxonomic lineages for MAGs.

## Methods

### Simulation of known truth dataset

The composition of a given environmental microbiome sample is of course unknown, and so it is difficult to measure classification accuracy on metagenomic data. Instead, data of known composition (“ground truth data”), such as simulated datasets or mock communities [46] are typically used to assess accuracy.

Here, InSilicoSeq (version 1.4.6) was used to generate simulated metagenomic data: 50 million paired-end reads using the HiSeq model with an exponential distribution [47] from known sequences. The input genomes used to create the data were 460 publicly available bacterial and archaeal reference genomes from the Hungate collection [10]. Since some of the Hungate collection are multi-contig, they were treated as draft genomes during data generation, using the *--draft* option. Complete genomes with a single contig were treated as such, using the *--genomes* option. A list of the Hungate genome files, and which are single or multi-contig, can be found in Supplementary Table S3.

As the simulated reads originated from the Hungate genomes, each read had a corresponding genome and therefore corresponding taxonomy. In this study the simulated data is referred to as “ground truth”, as the true taxonomy of each read is known. The number of reads simulated from each genome, and therefore for each taxonomy, were determined (using Ete3 [48]). The number of reads produced for each genome provided the number of reads produced for each taxon at the phylum, family, genus and species levels. This “ground truth” information was used to assess the classification accuracy of each read (see Figures 3 and 4, and Supplementary Figure S1 and Supplementary Tables S1 and S2).

### Design, choice and creation of reference databases

Six reference databases were used to classify the simulated metagenome, the details of which can be seen in Table 1. Each database was built using NCBI taxonomy downloaded on 07/03/2020. NCBI libraries for the RefSeq database were downloaded on 24/03/2020.

The Hungate reference database contains genomes from 460 rumen-dwelling microbes cultured in the Hungate 1000 project. These were the same genomes that were used to create the simulated metagenome; therefore, this database was fully representative of the data being classified. The Hungate database therefore acted as the ‘best case’ scenario for database choice, and can be seen as a positive control, as each read from the simulated metagenome should be represented in the Hungate database.

The RefSeq database is the standard Kraken2 [30] reference database (see [49]) widely used for taxonomy classification. It contains the complete collection of genomes in RefSeq for bacterial, archaeal and viral domains, the human genome and a collection of vectors (UniVec_core).

The Mini reference database is also a popular database for Kraken2 users, designed for users with low-memory computing environments. Both the Standard and Mini databases contain the same RefSeq reference genomes, but the Mini database was built using a hash function to down-sample minimisers, as described in the Kraken 2 manual and shown in Table 1 (*--max-db-size function*). The hash file for the Standard Kraken 2 database is 43 GB, whereas it is only 7.5 GB for the Mini Kraken 2 database. As this database is significantly smaller than the Standard reference database, read classification requires less memory. As the Mini reference database may be the first choice for users with limited computational resources, it was included in this study.

The RUG reference database contains 4,941 rumen MAGs assembled by Stewart *et al*. [17]. Whilst different from the cultured Hungate genomes, these assembled genomes were assembled from metagenomes also originating in the rumen. This custom database was included in the study to investigate the impact of a reference database containing assembled genomes on taxonomic classification.

The RefRUG and RefHun reference databases contain the complete collection of genomes in RefSeq (bacterial, viral and archaeal domains, the human genome and UniVec_Core vectors) in addition to the RUGs and Hungate genomes, respectively. These were included to investigate whether adding genomes or draft genomes from the same type of environmental microbiota as the data being classified improves taxonomic classification.

### Read classification using Kraken2

The simulated metagenome was classified using Kraken2 (version 2.0.8_beta) with the six reference databases described above. Default settings were used with the *--paired* option to accommodate the paired-end reads of the simulated metagenome.

Classification status was extracted from the Kraken output files and used to assign reads to one of two classes: classified or unclassified. The taxonomic ID for each read was extracted from the Kraken output files, and classified reads were compared to their known ground truth at the species, genus, family and phylum level (using Ete3). The reads were firstly grouped into “correct” or “incorrect” and then subsequently into “correct”, “incorrect”, “unclassified at this level”, “unclassified at any level” and “truth unknown”.

Finally, the Kraken 2 report files were used to compare read classification counts for each taxonomic level against the ground truth, and R^2^ calculated as the sum-of-squares of absolute deviation from the ground-truth.

## Declarations

### Ethics approval and consent to participate

Not applicable

### Consent for publication

Not applicable

### Competing interests

The authors declare that they have no completing interests

### Funding

The Roslin Institute forms part of the Royal (Dick) School of Veterinary Studies, University of Edinburgh. This project was supported by the Biotechnology and Biological Sciences Research Council (BBSRC; BB/S006680/1, BB/R015023/1), including institute strategic program grant BBS/E/D/30002276. R.H.S. is supported by an EASTBIO studentship funded by BBSRC (BB/M010996/1). A.W.W. and the Rowett Institute receive core financial support from the Scottish Government Rural and Environmental Sciences and Analytical Services (SG-RESAS).

### Author’s contributions

R.H.S. created the simulated data, conducted data analyses and bioinformatics, made figures, and contributed to writing the manuscript. M.W. conceived the study, carried out bioinformatics work and created figures. M.W., A.W.W. and L.G. supervised the project and contributed to writing the manuscript. All authors read and approved the final manuscript.

## Acknowledgements

We would like to thank all of those who were involved in creating and publicly sharing both the Hungate Collection data and the RUG data.

## Notes

### Competing Interest Statement

The authors have declared no competing interest.

## References

1. Kamra DN. Rumen microbial ecosystem. Curr Sci. 2005;89:124–35.

2. Auffret MD, Stewart RD, Dewhurst RJ, Duthie CA, Watson M, Roehe R. Identification of microbial genetic capacities and potential mechanisms within the rumen microbiome explaining differences in beef cattle feed efficiency. Front Microbiol. 2020;11 June:1–16.

3. Huws SA, Creevey CJ, Oyama LB, Mizrahi I, Denman SE, Popova M, et al. Addressing global ruminant agricultural challenges through understanding the rumen microbiome: past, present, and future. Front Microbiol. 2018;9:1–33.

4. Martínez-Álvaro M, Auffret MD, Stewart RD, Dewhurst RJ, Duthie CA, Rooke JA, et al. Identification of complex rumen microbiome interaction within diverse functional niches as mechanisms affecting the variation of methane emissions in bovine. Front Microbiol. 2020;11:1–13.

5. Roehe R, Dewhurst RJ, Duthie CA, Rooke JA, McKain N, Ross DW, et al. Bovine host genetic variation influences rumen microbial methane production with best selection criterion for low methane emitting and efficiently feed converting hosts based on metagenomic gene abundance. PLoS Genet. 2016;12:1–20.

6. Wallace RJ, Rooke JA, McKain N, Duthie CA, Hyslop JJ, Ross DW, et al. The rumen microbial metagenome associated with high methane production in cattle. BMC Genomics. 2015;16:1–14.

7. Auffret MD, Stewart R, Dewhurst RJ, Duthie CA, Rooke JA, Wallace RJ, et al. Identification, comparison, and validation of robust rumen microbial biomarkers for methane emissions using diverse Bos Taurus breeds and basal diets. Front Microbiol. 2018;8:1–15.

8. Auffret MD, Dewhurst RJ, Duthie CA, Rooke JA, John Wallace R, Freeman TC, et al. The rumen microbiome as a reservoir of antimicrobial resistance and pathogenicity genes is directly affected by diet in beef cattle. Microbiome. 2017;5:1–11.

9. Henderson G, Cox F, Ganesh S, Jonker A, Young W, Janssen PH, et al. Rumen microbial community composition varies with diet and host, but a core microbiome is found across a wide geographical range. Sci Rep. 2015;5.

10. Seshadri R, Leahy SC, Attwood GT, Teh KH, Lambie SC, Cookson AL, et al. Cultivation and sequencing of rumen microbiome members from the Hungate1000 Collection. Nat Biotechnol. 2018;36:359–67.

11. Creevey CJ, Kelly WJ, Henderson G, Leahy SC. Determining the culturability of the rumen bacterial microbiome. Microb Biotechnol. 2014;7:467–79.

12. Quince C, Walker AW, Simpson JT, Loman NJ, Segata N. Shotgun metagenomics, from sampling to analysis. Nat Biotechnol. 2017;35:833–44.

13. Stewart RD, Auffret MD, Warr A, Wiser AH, Press MO, Langford KW, et al. Assembly of 913 microbial genomes from metagenomic sequencing of the cow rumen Robert. Nat Commun. 2018;9:1–11.

14. Rappé MS, Giovannoni SJ. The uncultured microbial majority. Annu Rev Microbiol. 2003;57:369–94.

15. Lewis WH, Tahon G, Geesink P, Sousa DZ, Ettema TJG. Innovations to culturing the uncultured microbial majority. Nat Rev Microbiol. 2021;19:225–40.

16. Watson M. New insights from 33,813 publicly available metagenome-assembled-genomes (MAGs) assembled from the rumen microbiome. Preprint at https://www.biorxiv.org/content/10.1101/2021.04.02.438222v1.full (2021).

17. Stewart RD, Auffret MD, Warr A, Walker AW, Roehe R, Watson M. Compendium of 4,941 rumen metagenome-assembled genomes for rumen microbiome biology and enzyme discovery. Nat Biotechnol. 2019;37:953–61.

18. Solden LM, Naas AE, Roux S, Daly RA, Collins WB, Nicora CD, et al. Interspecies cross-feeding orchestrates carbon degradation in the rumen ecosystem. Nat Microbiol. 2018;3:1274–84.

19. Glendinning L, Genç B, Wallace RJ, Watson M. Metagenomic analysis of the cow, sheep, reindeer and red deer rumen. Sci Rep. 2021;11:3–12.

20. Wilkinson T, Korir D, Ogugo M, Stewart RD, Watson M, Paxton E, et al. 1200 high-quality metagenome-assembled genomes from the rumen of African cattle and their relevance in the context of sub-optimal feeding. Genome Biol. 2020;21:1–25.

21. Parks DH, Rinke C, Chuvochina M, Chaumeil PA, Woodcroft BJ, Evans PN, et al. Recovery of nearly 8,000 metagenome-assembled genomes substantially expands the tree of life. Nat Microbiol. 2017;2:1533–42.

22. Xie F, Jin W, Si H, Yuan Y, Tao Y, Liu J, et al. An integrated gene catalog and over 10,000 metagenome-assembled genomes from the gastrointestinal microbiome of ruminants. Microbiome. 2021;9:1–20.

23. Svartström O, Alneberg J, Terrapon N, Lombard V, De Bruijn I, Malmsten J, et al. Ninety-nine de novo assembled genomes from the moose (Alces alces) rumen microbiome provide new insights into microbial plant biomass degradation. ISME J. 2017;11:2538–51.

24. Glendinning L, Stewart RD, Pallen MJ, Watson KA, Watson M. Assembly of hundreds of novel bacterial genomes from the chicken caecum. Genome Biol. 2020;21:1–16.

25. Peng X, Wilken SE, Lankiewicz TS, Gilmore SP, Brown JL, Henske JK, et al. Genomic and functional analyses of fungal and bacterial consortia that enable lignocellulose breakdown in goat gut microbiomes. Nat Microbiol. 2021;6:499–511.

26. Li J, Zhong H, Ramayo-Caldas Y, Terrapon N, Lombard V, Potocki-Veronese G, et al. A catalog of microbial genes from the bovine rumen unveils a specialized and diverse biomass-degrading environment. Gigascience. 2020;9:1–15.

27. Hess M, Sczyrba A, Egan R, Kim TW, Chokhawala H, Schroth G, et al. Metagenomic discovery of biomass-degrading genes and genomes from cow rumen. Science. 2011;331:463–7.

28. Gharechahi J, Vahidi MF, Bahram M, Han JL, Ding XZ, Salekdeh GH. Metagenomic analysis reveals a dynamic microbiome with diversified adaptive functions to utilize high lignocellulosic forages in the cattle rumen. ISME J. 2021;15:1108–20.

29. Wood DE, Salzberg SL. Kraken: Ultrafast metagenomic sequence classification using exact alignments. Genome Biol. 2014.

30. Wood DE, Lu J, Langmead B. Improved metagenomic analysis with Kraken 2. Genome Biol. 2019;20:1–13.

31. Blackwell GA, Hunt M, Malone KM, Lima L, Horesh G, Alako BTF, et al. Exploring bacterial diversity via a curated and searchable snapshot of archived DNA sequences. PLoS Biol. 2021;19.

32. Méric G, Wick RR, Watts SC, Holt KE, Inouye M. Correcting index databases improves metagenomic studies. Preprint at https://www.biorxiv.org/content/10.1101/712166v1 (2019).

33. O’Leary NA, Wright MW, Brister JR, Ciufo S, Haddad D, McVeigh R, et al. Reference sequence (RefSeq) database at NCBI: current status, taxonomic expansion, and functional annotation. Nucleic Acids Res. 2016;44:D733–45.

34. Nasko DJ, Koren S, Phillippy AM, Treangen TJ. RefSeq database growth influences the accuracy of k-mer-based lowest common ancestor species identification. Genome Biol. 2018;19:1–10.

35. R. Marcelino V, Holmes EC, Sorrell TC. The use of taxon-specific reference databases compromises metagenomic classification. BMC Genomics. 2020;21:1–5.

36. McIntyre ABR, Ounit R, Afshinnekoo E, Prill RJ, Hénaff E, Alexander N, et al. Comprehensive benchmarking and ensemble approaches for metagenomic classifiers. Genome Biol. 2017;18:1–19.

37. Pallen MJ, Telatin A, Oren A. The Next Million Names for Archaea and Bacteria. Trends Microbiol. 2021;29:289–98.

38. Walker AW. Microbiota of the Human Body. 2016;902:5–32.

39. Schoch CL, Ciufo S, Domrachev M, Hotton CL, Kannan S, Khovanskaya R, et al. NCBI taxonomy: a comprehensive update on curation, resources and tools. Database. 2020;2020:1–21.

40. Breitwieser FP, Lu J, Salzberg SL. A review of methods and databases for metagenomic classification and assembly. Brief Bioinform. 2018;20:1125–39.

41. Murray AE, Freudenstein J, Gribaldo S, Hatzenpichler R, Hugenholtz P, Kämpfer P, et al. Roadmap for naming uncultivated Archaea and Bacteria. Nat Microbiol. 2020.

42. Collins MD, Lawson PA, Willems A, Cordoba JJ, Fernandez-Garayzabal J, Garcia P, et al. The phylogeny of the genus Clostridium: proposal of five new genera and eleven new species combinations. Int J Syst Bacteriol. 1994;44:812–26.

43. Parks DH, Chuvochina M, Waite DW, Rinke C, Skarshewski A, Chaumeil PA, et al. A standardized bacterial taxonomy based on genome phylogeny substantially revises the tree of life. Nat Biotechnol. 2018;36:996.

44. Thompson CC, Amaral GR, Campeão M, Edwards RA, Polz MF, Dutilh BE, et al. Microbial taxonomy in the post-genomic era: rebuilding from scratch? Arch Microbiol. 2015;197:359–70.

45. Parks DH, Chuvochina M, Chaumeil PA, Rinke C, Mussig AJ, Hugenholtz P. A complete domain-to-species taxonomy for Bacteria and Archaea. Nat Biotechnol. 2020.

46. Bokulich NA, Rideout JR, Mercurio WG, Shiffer A, Wolfe B, Maurice CF, et al. mockrobiota: a Public Resource for Microbiome Bioinformatics Benchmarking. mSystems. 2016;1.

47. Gourlé H, Karlsson-Lindsjö O, Hayer J, Bongcam-Rudloff E. Simulating Illumina metagenomic data with InSilicoSeq. Bioinformatics. 2019.

48. Huerta-Cepas J, Serra F, Bork P. ETE 3: reconstruction, analysis, and visualization of phylogenomic data. Mol Biol Evol. 2016;33:1635–8.

49. Wood DE. Kraken 2 Standard Reference Database. https://github.com/DerrickWood/kraken2/wiki/Manual#standard-kraken-2-database. Accessed 16 Mar 2020.

