## Additional file 1 for "Investigating the impact of database choice on the accuracy of metagenomic read classification for the rumen microbiome"

**Supplementary Table S1** *Classification status of reads for the six reference databases at various taxonomic levels.*

| | Overall | | Phylum | | Family | | Genus | | Species | |
|---|---|---|---|---|---|---|---|---|---|---|
| Database | Classified | Unclassified | Classified | Unclassified | Classified | Unclassified | Classified | Unclassified | Classified | Unclassified |
| Hungate | 99.95 | 0.05 | 99.94 | 0.06 | 97.99 | 2.01 | 82.58 | 17.42 | 92.69 | 7.31 |
| Mini | 39.85 | 60.15 | 39.40 | 60.60 | 38.44 | 61.56 | 37.21 | 62.79 | 32.20 | 67.80 |
| RefSeq | 50.28 | 49.72 | 49.62 | 50.38 | 48.26 | 51.74 | 46.77 | 53.23 | 43.27 | 56.73 |
| RUG | 45.66 | 54.34 | 45.16 | 54.84 | 42.36 | 57.64 | 27.99 | 72.01 | 43.93 | 56.07 |
| RefRUG | 70.09 | 29.91 | 69.53 | 30.47 | 67.05 | 32.95 | 54.69 | 45.31 | 61.53 | 38.47 |
| RefHun | 99.96 | 0.04 | 99.93 | 0.07 | 97.84 | 2.16 | 82.13 | 17.87 | 89.27 | 10.73 |

Classification status refers to whether the read was classified, or unclassified, regardless of accuracy. Each row denotes the six databases used to classify reads with Kraken2. The “Overall” column refers to the percentage of reads which were classified or unclassified by Kraken2 regardless of taxonomic level. Subsequent columns refer to the percentage of reads which were classified or unclassified by Kraken2 at various taxonomic levels as shown in the column headers.

**Supplementary Table S2** *Classification status of reads compared to the ground truth for the six reference databases at various taxonomic levels.*

| | Status | Phylum | Family | Genus | Species |
|---|---|---|---|---|---|
| Hungate | Correct | 99.94 | 97.99 | 82.56 | 92.56 |
| | Incorrect | 0.00 | 0.00 | 0.01 | 0.13 |
| | Truth unknown | 0.00 | 1.86 | 16.32 | 0.00 |
| | Unclassified at any level | 0.05 | 0.05 | 0.05 | 0.05 |
| | Unclassified at this level | 0.01 | 0.10 | 1.07 | 7.26 |
| Mini | Correct | 38.24 | 35.62 | 32.04 | 20.65 |
| | Incorrect | 1.16 | 2.74 | 3.82 | 11.55 |
| | Truth unknown | 0.00 | 1.86 | 16.32 | 0.00 |
| | Unclassified at any level | 60.15 | 58.55 | 45.74 | 60.15 |
| | Unclassified at this level | 0.45 | 1.23 | 2.08 | 7.65 |
| RefHun | Correct | 99.92 | 97.82 | 81.90 | 88.92 |
| | Incorrect | 0.01 | 0.01 | 0.23 | 0.35 |
| | Truth unknown | 0.00 | 1.86 | 16.32 | 0.00 |
| | Unclassified at any level | 0.04 | 0.04 | 0.04 | 0.04 |
| | Unclassified at this level | 0.03 | 0.27 | 1.52 | 10.69 |
| RefRUG | Correct | 67.43 | 59.00 | 47.31 | 25.87 |
| | Incorrect | 2.09 | 7.13 | 5.11 | 35.65 |
| | Truth unknown | 0.00 | 1.86 | 16.32 | 0.00 |
| | Unclassified at any level | 29.91 | 29.51 | 21.63 | 29.91 |
| | Unclassified at this level | 0.56 | 2.51 | 9.63 | 8.56 |
| RefSeq | Correct | 46.13 | 40.93 | 35.97 | 22.74 |
| | Incorrect | 3.49 | 7.07 | 7.85 | 20.53 |
| | Truth unknown | 0.00 | 1.86 | 16.32 | 0.00 |
| | Unclassified at any level | 49.72 | 48.34 | 37.22 | 49.72 |
| | Unclassified at this level | 0.65 | 1.81 | 2.65 | 7.01 |
| RUG | Correct | 43.69 | 35.76 | 26.11 | 8.46 |
| | Incorrect | 1.47 | 5.71 | 1.29 | 35.47 |
| | Truth unknown | 0.00 | 1.86 | 16.32 | 0.00 |
| | Unclassified at any level | 54.35 | 53.80 | 44.41 | 54.35 |
| | Unclassified at this level | 0.49 | 2.87 | 11.88 | 1.72 |

The databases and detailed classification status are shown in the first column. Subsequent columns contain the percentage of reads at that taxonomic level, which had been classified by the database and had the particular classification status outlined in the first column. “Correct” and “incorrect” refer to reads which were classified correctly or incorrectly by Kraken2 using the respective database. “Truth unknown” refers to the reads that originate from genomes that do not have an assigned family or genus. “Unclassified at any level” refers to reads that were not classified to any taxonomic level. “Unclassified at this level” refers to reads which were classified at other taxonomic levels, but not the level being examined in a given column.

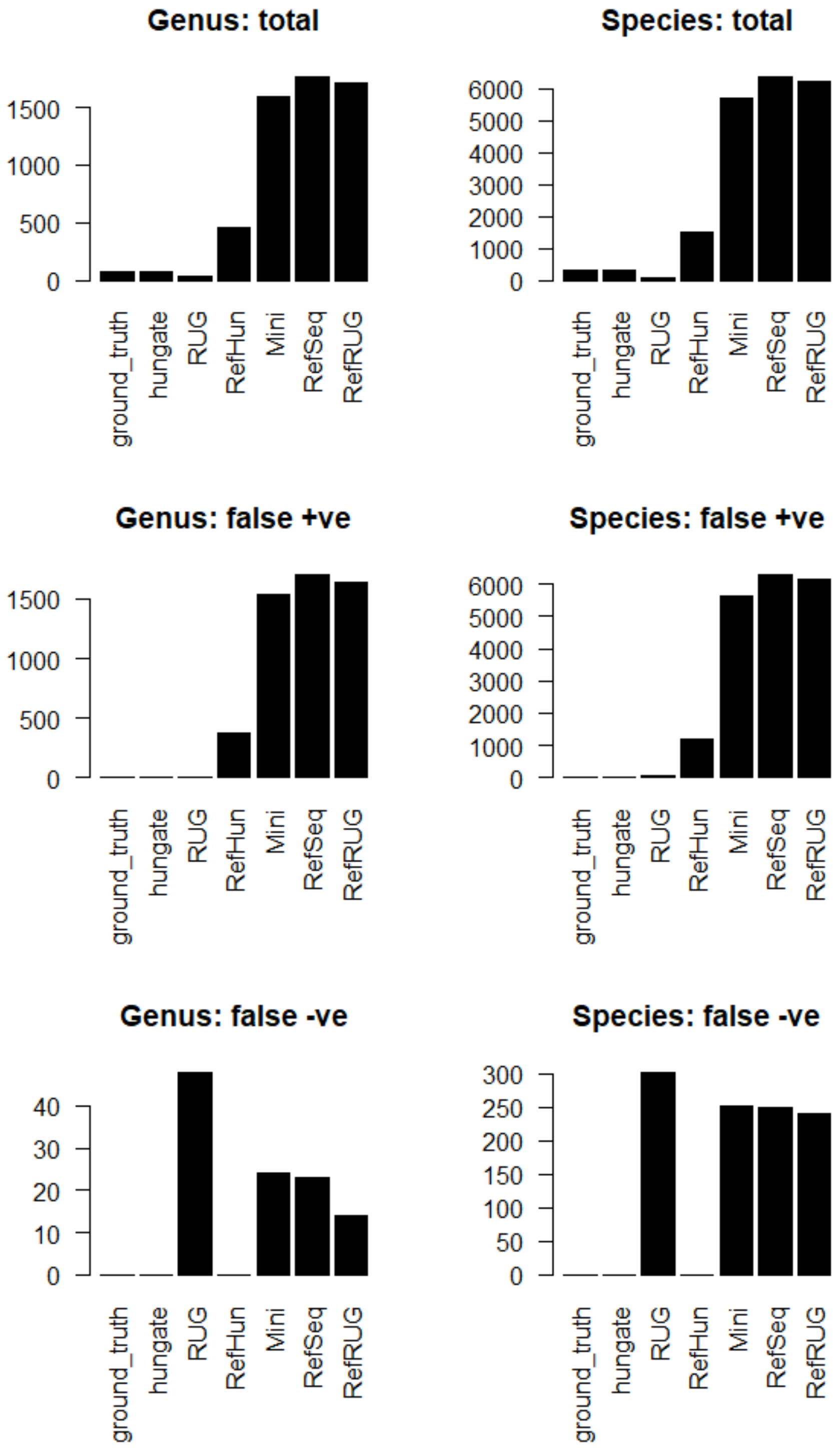


**Supplementary Figure S1** The frequency of genera and species in the ground truth data, and in the classification results for each reference database. The total frequency is shown in the top two graphs, the middle graphs show the frequency of false positives occurring, and the bottom two graphs show the frequency of false negatives.
